# Life in coastal pebble sediment: Unique interstitial organism community and selective feeding on meiobenthos by interstitial fishes (*Luciogobius*: Gobiidae)

**DOI:** 10.1101/499194

**Authors:** Kasumi Kondo, Makoto Kato

## Abstract

Interstitial areas of coastal pebble sediment in the Japanese Archipelago are inhabited by extremely elongated gobies of the genus *Luciogobius*, which are characterized by an increased number of vertebrae and reduction of scales, eyes, and fins. To explore the little-known interstitial life of *Luciogobius* gobies, we investigated the diets of two interstitial *Luciogobius* species, *L. elongatus* and *L. grandis*, and the interstitial organism communities of the gobies’ microhabitats in an exposed gravelly coast in Shirahama, southern Japan. The interstitial organism community in pebbly sediment was dominated by minute arthropods such as harpacticoids, isopods, and ostracods, presenting a marked contrast to the communities in sandy sediments, which are dominated by nematodes and turbellarians. The gut contents of the two goby species were composed exclusively of interstitial organisms, especially harpacticoides and isopods. Although each prey assemblage was roughly similar to the interstitial organism community in the corresponding microhabitat, marked preferences for harpacticoids and flabelliferan isopods were detected in *L. elongatus* and *L. grandis*, respectively. Irrespective of their intense feeding of harpacticoids, rare catches of large isopods were suggested to be nutritionally important for the gobies. These results suggest that the *Luciogobius* gobies are the first known fishes that depend exclusively on interstitial organisms, and that selective feeding upon meiobenthos may facilitate the coexistence of several interstitial goby species in pebbly sediment.

## Introduction

The Japanese Archipelago is located on a tectonic plate boundary and contains steep mountain ranges shaped by active tectonic uplift [1]. The rough landscape and heavy rainfall caused by monsoons have contributed to the formation of many gravelly beaches along the sea coasts. The substrata of these gravelly beaches are mainly composed of pebbles, and diverse and numerous interstitial invertebrates inhabit the interstitial spaces among pebbles [2]. Because the pebbles are invariably stirred by waves, the interstitial habitats seem to be too dynamic for vertebrates to inhabit. However, the gravelly beaches in the Japanese Archipelago are inhabited by diverse *Luciogobius* gobies, which have flexible, elongated bodies [3].

The genus *Luciogobius* comprises 17 described and more than 20 undescribed species, and its members are characterized by a slender, extremely elongated body with highly reduced eyes and fins. This genus has experienced an adaptive radiation in gravelly or rocky coasts around the Japanese Archipelago [3–4]. While *Luciogobius* species are distributed in East Asia, ranging from Japan to the Korean Peninsula, southernmost seacoasts of Russia, Taiwan, Hong Kong, and Hainan Island [4–5], its diversity is concentrated in the Japanese Archipelago.

Among *Luciogobius* species, a small number are lapidicolous (living under stones) or cavicolous (living in caves), while others inhabit pebbly interstitial habitats in gravelly coasts. These interstitial *Luciogobius* species are remarkably diverse, because there are many local endemic species, and because several *Luciogobius* species live sympatrically. Thus, it is intriguing to consider why rapid diversification has occurred in within fish lineages in interstitial habitats on gravelly beaches. Preliminary observations have shown that sympatric *Luciogobius* species segregate their microhabitats by preferring sediment layers with specific pebble size distributions [3].

Because most goby species are carnivores of benthic organisms, differences in microhabitats among species must cause differences in potential prey items. Among *Luciogobius* species, diet has been reported only in a lapidicolous species, *Luciogobius guttatus* [6–7], which feeds mainly on small benthic arthropods (e.g., crabs, shrimps, hermit crabs, amphipods, and harpacticoids). To explore adaptive radiation among *Luciogobius* species, we need to clarify both the diets of the diverse interstitial *Luciogobius* species and the potential prey organisms living in the interstitial microhabitats.

Sediment on sandy beaches harbors diverse and numerous minute interstitial organisms, such as foraminiferans, nematodes, annelids, harpacticoids, and ostracods, and these organisms have adapted to the dynamic interstitial life by having flexible, elastic, extended, or armored bodies [8]. In contrast with the accumulated information on interstitial organisms from sandy beaches, information on interstitial organisms in pebbly sediment is scarce. It is noteworthy that interstitial vertebrates inhabit pebbly but never sandy interstitial environments. Thus, to understand interstitial life in pebbly sediment we must study the community structure of interstitial organisms in pebbly habitats.

To explore adaptive radiation of gobies in pebbly sediment, we focused on two interstitial *Luciogobius* species: one extremely thin and one thicker species. We investigated the fishes’ diets, as well as the communities of potential prey organisms for these species, both of which inhabit interstitial sediment of gravelly beaches in the Japanese Archipelago. These two species are sympatric at a beach scale, but segregate their microhabitats. This is the first study reporting the diet of interstitial fishes, and simultaneously the first study reporting on the communities of interstitial organisms in pebbly sediment of gravelly coasts.

## Materials and methods

### Ethics statement

The field sampling and sample treatment were conducted in accordance with the “Guidelines for the use of fishes in research” by the Ichthyological Society of Japan (http://www.fish-isi.ip/english/guidelines.html). All animal experiments were approved by the Ethics Committee for Animal Experiments of Kyoto University. The experimental procedures were conducted in accordance with the approved guidelines.

### Study site

This study was conducted at a gravelly beach (33°41’39.8”N, 135°20’08.3”E) near the Seto Marine Biological Laboratory of Kyoto University in Shirahama, Wakayama prefecture, Japan, where the maximum tidal range is about 200 cm. The beach was slightly protected from strong waves by a cape facing the Pacific and composed of pebbly sediment washed by strong waves during high tide. The intertidal zones of the pebbly sediment are inhabited by five *Luciogobius* species, two of which are common: *L. elongatus* and *L. grandis*. No specific permissions were required for these locations, and neither endangered nor protected species were involved in this field study.

### Sampling of gobies

Two *Luciogobius* species, *L. elongatus* and *L. grandis*, were sampled from intertidal zones of the beach by digging pebbly sediment with a shovel (Figs 1, 2). The two goby species live on the same gravelly beach, but their microhabitats differ. Namely, the sediment was coarser in *L. grandis* habitats than in *L. elongatus* habitats. *Luciogobius* gobies were found mainly in the middle or lower layers of the sediment piled on bedrock. Sampling was carried out during the daytime and nighttime, during low tide during the spring tide, from 24 to 26 February, 11 to 13 June, and 4 to 6 October 2017. To examine gut contents, all collected gobies were immediately fixed in 10% formalin.

**Fig 1.**
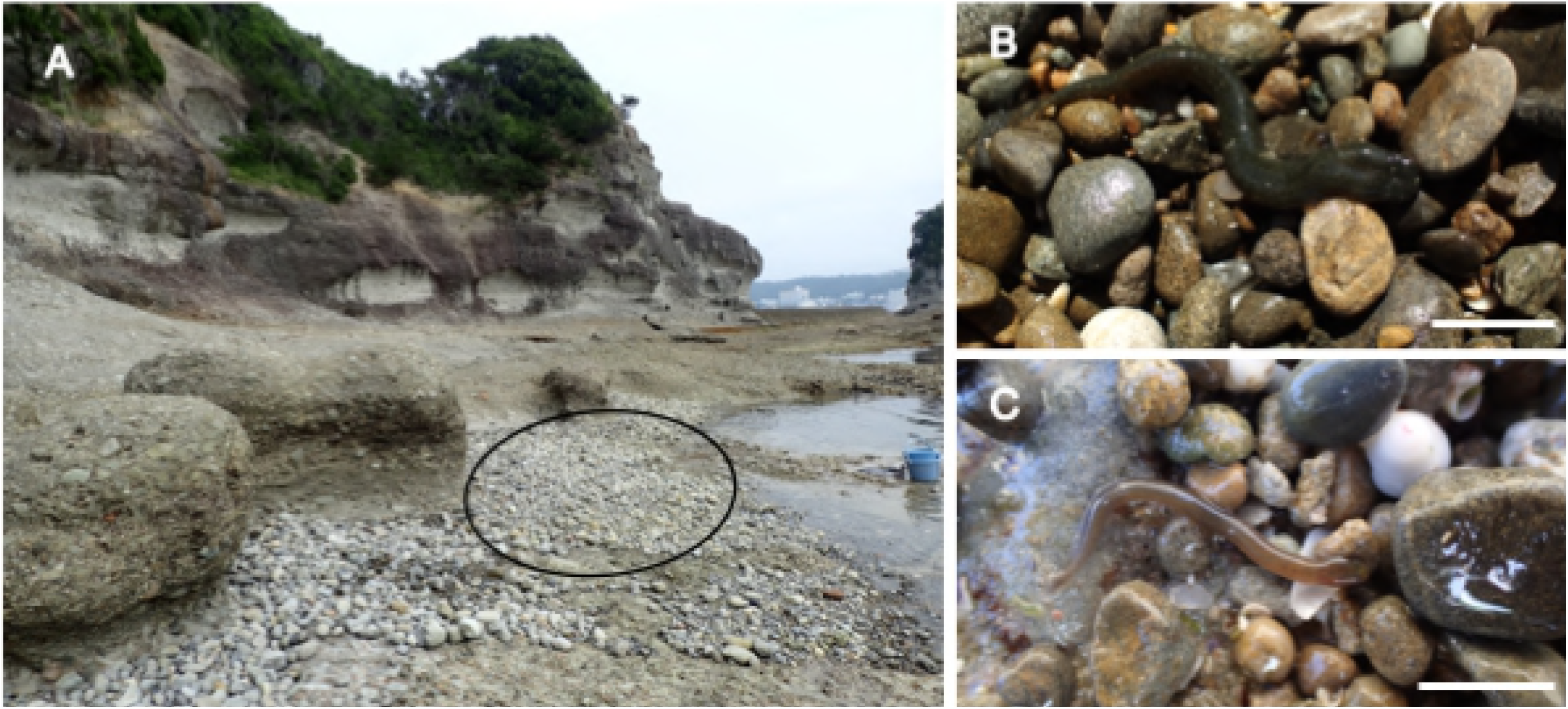
(A) Landscape of the study site and live (B) *Luciogobius grandis* and (C) *L. elongatus* gobies in their microhabitats at Shirahama. *Scale bar* = 1 cm.

**Fig 2.**
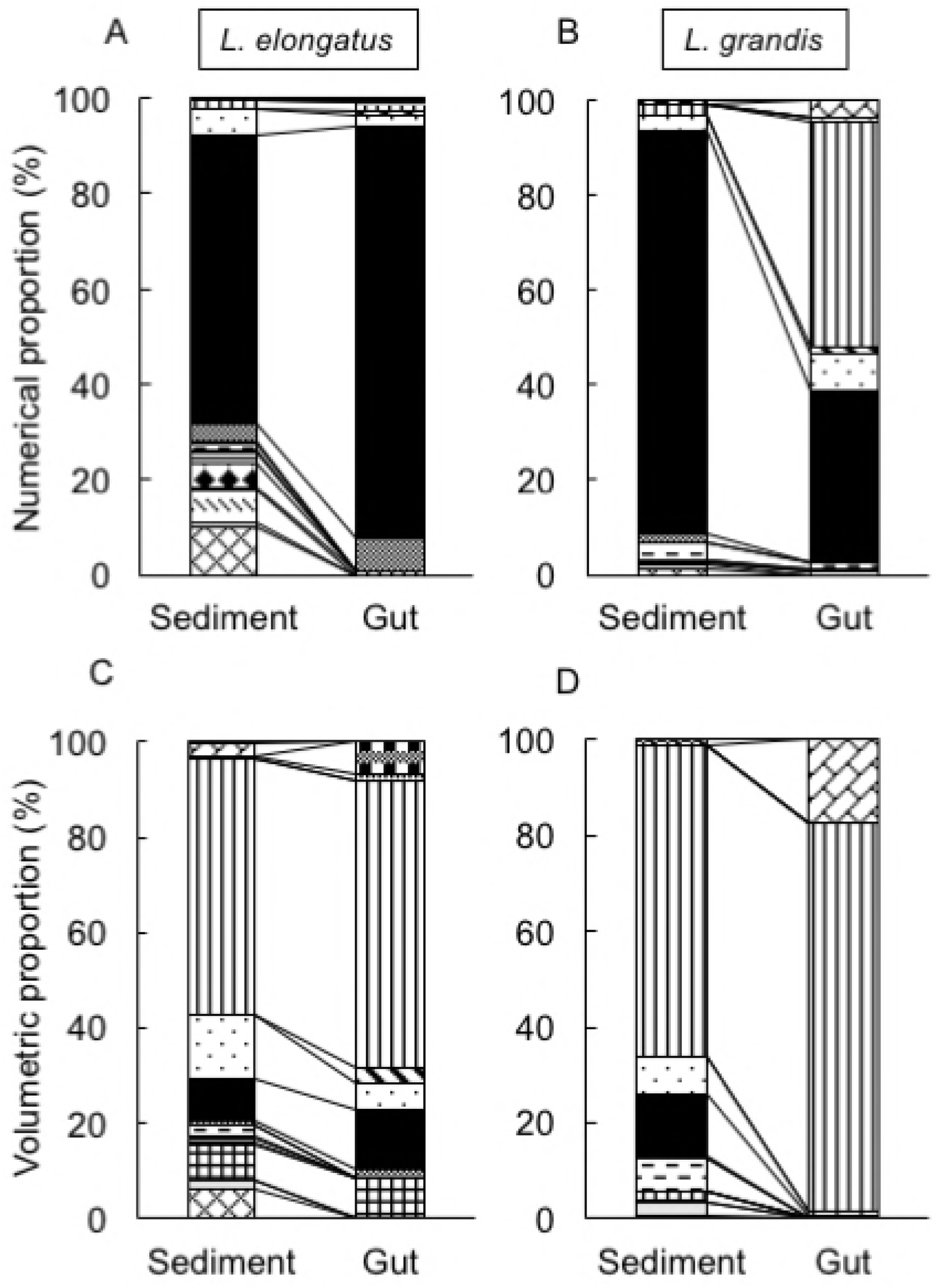
Photographs (A) *Luciogobius elongatus* and (B) *L. grandis*. *Scale bar* = 1 cm.

### Gut content

After removing formalin, we dissected the stomach of each goby, extracted all bodies and fragments of prey individuals, sorted them into taxonomic groups, and counted them under a binocular microscope. Thus, we obtained prey assemblage data sets for two goby species (*L. elongatus* and *L. grandis*), for three seasons (February, June, and October), and for daytime and nighttime. Based on the datasets, we calculated the percentage of fish that ate at least one individual of the prey group (%F). Furthermore, we estimated the numeric proportion of each prey group (%N), i.e., the numeric percentage of each prey item out of the total number of individuals. To detect factors involving in interspecific, seasonal and diurnal variances of the prey assemblages, we conducted nonmetric multidimensional scaling (NMDS) using Soft [9] and obtained a two-dimensional graphical representation of the multivariate prey assemblages.

### Interstitial organism community in goby microhabitats

To uncover the community structure of potential prey organisms, we sampled interstitial organisms in the gobies’ microhabitats, since preliminary observations suggested that their stomach contents were composed mainly of interstitial organisms. We sampled about 2 L of pebbly sediment from the same site as goby sampling, put it into a bucket, poured sea water into the bucket, stirred the water with a shovel for 30 s, and then filtered the supernatant through a plankton net (100-μm mesh). We repeated the filtering procedure three times. Collected interstitial organisms were immediately fixed in 5% formalin. In the laboratory, the interstitial organisms were dyed with Rose Bengal to more easily observe translucent bodies [8]. The samples were sorted into taxonomic groups in the same manner as gut contents and were counted.

To analyze sediment granularity, pebbly sediments from the gobies’ microhabitats were collected and sorted with standard sieves in running water. The sorted sediments were dried and weighed. From the data set of the fractions, the mean particle diameter and sorting index were calculated for each sediment sample.

### Estimation of prey biomass

To estimate the biomass of prey organisms, we measured the length (L) and width (W) of 10–20 individuals from each taxonomic group (if fewer than ten individuals were available, we measured as many individuals as possible), and calculated the average volume using the following formula [8]:

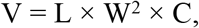

where C is the conversion factor given to each taxonomical group [8]. For groups for which the conversion factor had not been estimated, we approximated the individual’s body to a cylinder, a rectangular square, or another three-dimensional shape. Using this average volume data, we estimated the biomass compositions of both the prey assemblages and the interstitial organism communities.

## Results

### Microhabitats

We collected 52 *L. elongatus* and 75 *L. grandis* gobies from pebbly sediments in lower intertidal zones of a gravelly coast in Shirahama at spring low tide. Although both species lived sympatrically, their microhabitats differed. The mean sediment particle diameter was about 7 mm in both the *L. elongatus* and *L. grandis* microhabitats, but the sorting index was higher in *L. elongatus* microhabitats (0.54, i.e., moderately well-sorted) than *L. grandis* microhabitats (0.40, i.e., well-sorted).

### Seasonal patterns of growth and feeding

The mean total length of collected *L. elongatus* gobies was roughly 30 mm throughout the year (Fig 3), suggesting that most collected gobies were adults. In contrast, the mean total length of collected *L. grandis* gobies increased from February to October (Fig 3). In *L. grandis*, most gobies collected in October were adults, while gobies collected in February and June contained juveniles. In addition to gut contents, we monitored ovarian maturation of female gobies. *L. grandis* females had matured ovaries in October, while *L. elongatus* females had matured ovaries in June and October (Fig 4A).

**Fig 3.**
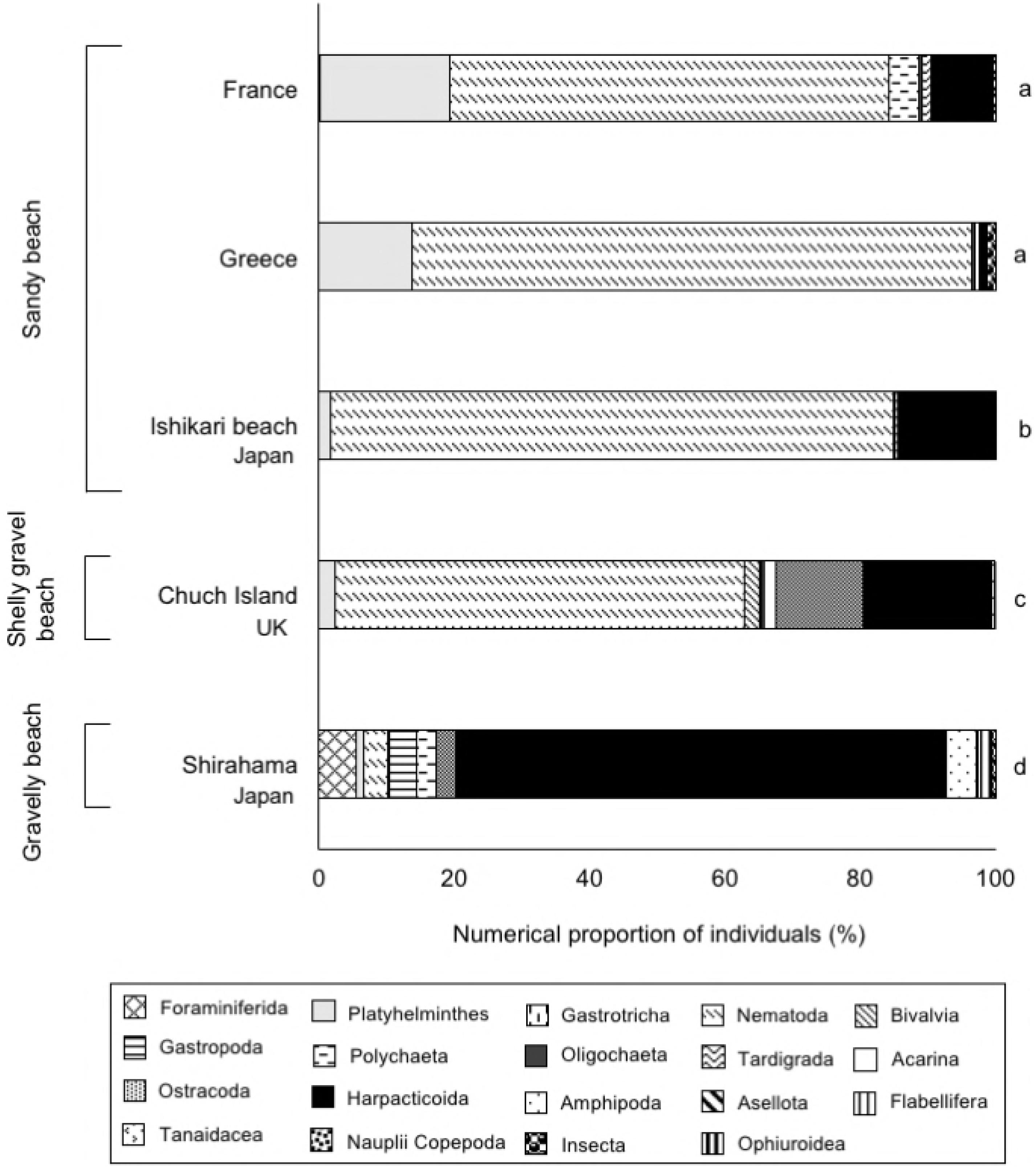
**Seasonal changes of seawater temperature (°C) and body lengths (mean ± standard deviation) of two species of gobies at Shirahama:** square, *Luciogobius elongatus*; diamond, *L. grandis*. Sampling of gobies was conducted three times at the dates shown by a–c.

**Fig 4.**
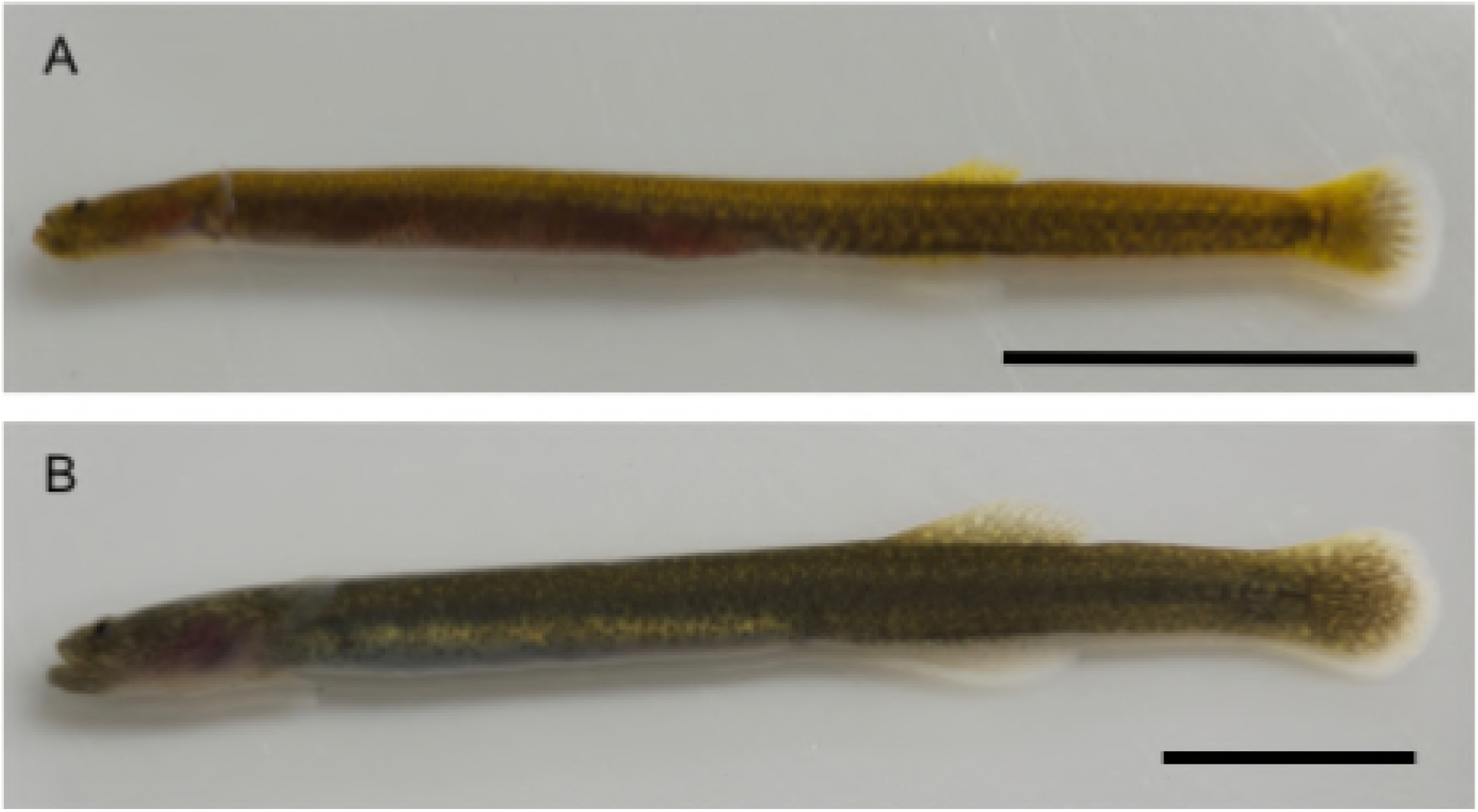
Seasonal changes in the percentage of (A) gravid females and (B) average number of prey individuals per gut.

The number of prey organisms in guts changed seasonally, and the mean was higher in *L. elongatus* than in *L. grandis* (Fig 4B). The interspecific difference in the number of prey organisms was greatest in June, when the gut was almost vacant in *L. grandis* but full in *L. elongatus*.

### Prey assemblages

Most organisms identified in gobies’ gut contents were less than 1 mm in length, while very few were 1–3 mm. Following the definition of meiobenthos [2], the main prey organisms of *Luciogobius* gobies were meiobenthos inhabiting pebbly sediment. The diets contained neither alga nor sand. The prey organisms comprised four phyla: Nematoda, Mollusca, Annelida, and Arthropoda (Table 1). The most common prey organisms were minute arthropods (Table 1), such as harpacticoids (Fig 5C), flabelliferan and asellotan isopods (Fig 5A, E), amphipods (Fig 5B), and ostracods (Fig 5D). The numerical proportion of arthropods in goby diets generally exceeded 95% in both species and reached 100% in the diurnal diets of *L. grandis* (Table 1). The prey assemblages in diet were largely similar between *L. elongatus* and *L. grandis*, but ostracods and an interstitial curviform gastropod, *Caecum glabella* (Caecidae: Rissoidea) (Fig 5G), were ingested only by *L. elongatus* (Table 1).

**Table 1.**
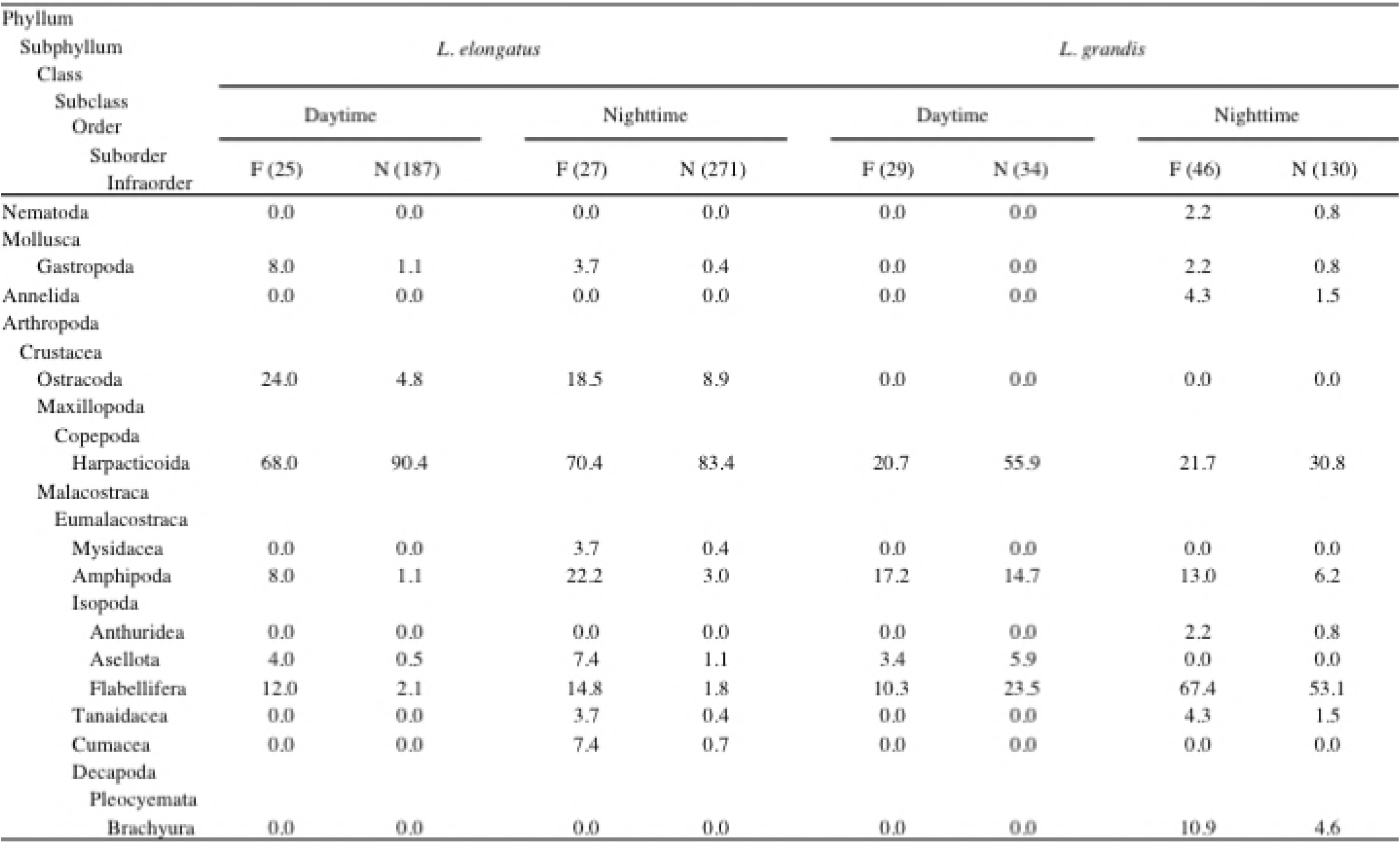
Prey organisms found in the guts of two goby species, *Luciogobius elongatus* and *L. grandis*, including their frequencies of occurrence and numerical proportions. Frequency of occurrence (F) is the proportion of gobies that contained at least one individual of each prey group in their gut (% gobies), and numerical proportion (N) is the proportion of each prey item out of all prey items in the gut (% prey individuals).

**Fig 5.**
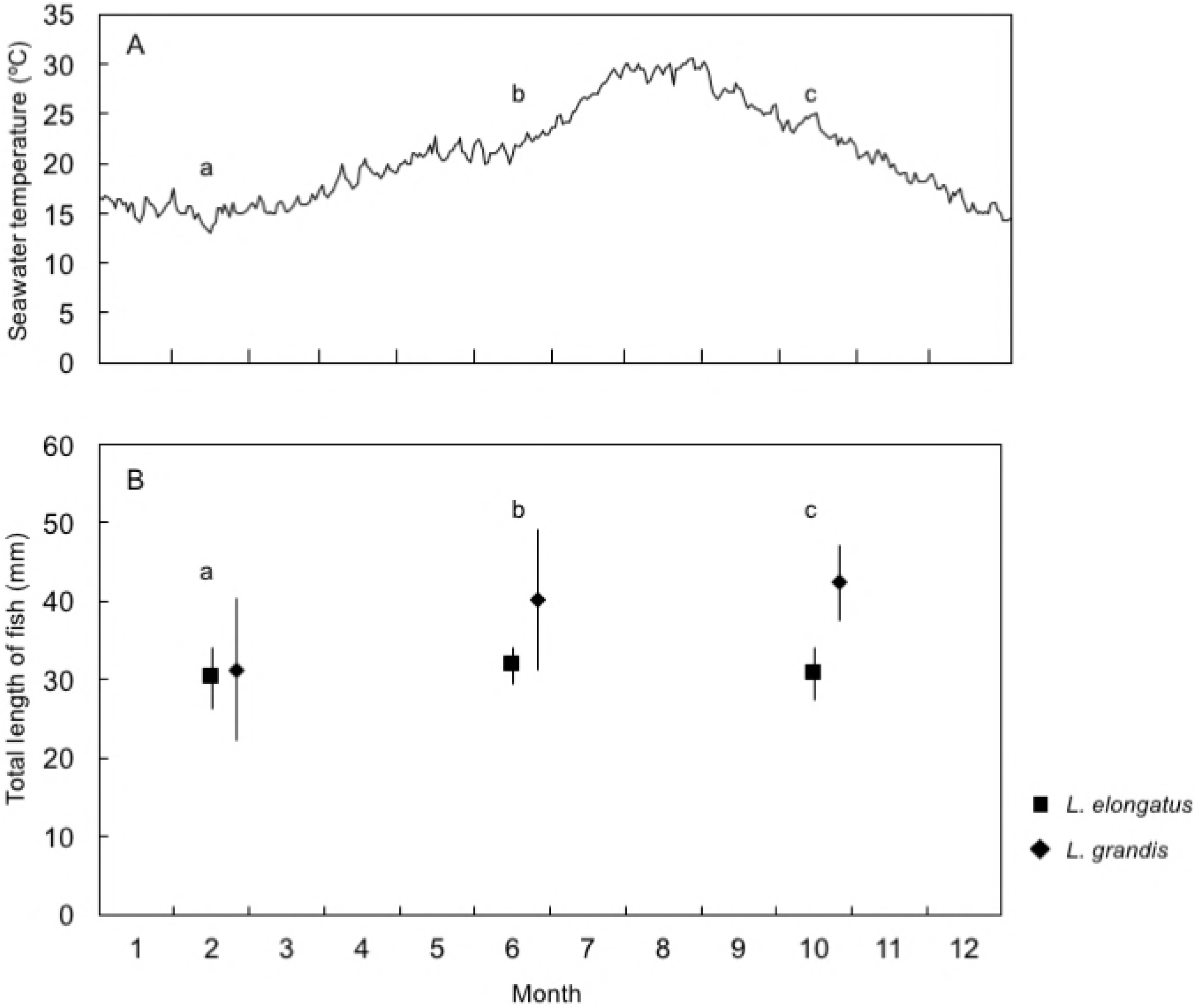
Photographs of the main interstitial meiobenthos groups collected from pebbly sediment in the gobies’ microhabitats: (A) flabelliferan isopod (Sphaeromatidae sp.); (B) amphipod; (C) harpacticoid copepod; (D) ostracod; (E) asellotan isopod (Janiridae sp.); (F) tanaid; (G) gastropod (*Caecum glabella*); (H) gastropod (*Ammonicera japonica*); (I) gastropod (Cingulopsidae sp.); (J) foraminiferans; (K) nematode; (L) insect (Collembola). These meiobenthos were dyed pink by Rose Bengal. *Scale bar* = 0.5 mm.

To compare prey assemblages between the two goby species, between the daytime and the nighttime and among three seasons (February, June, and October), NMDS analysis was conducted. NMDS ordination of the prey assemblages is shown in Fig 6, where the stress values were less than 0.2. The NMDS plots of *L. elongatus* and *L. grandis* suggested that the diet was not clearly discriminated between the two goby species, but that *L. elongatus* fed upon ostracods, cumaceans, and tanaidaceans more frequently than *L. grandis*, and that *L. grandis* fed upon flabelliferan isopods and brachyurans more frequently than *L. elongatus*. Harpacticoids and amphipods were common prey items for both gobies. There was no marked difference in diet between the daytime and the nighttime. The average number of prey items in each goby’s gut contents varied between the goby species, among seasons, and between daytime and nighttime (Fig 4B). *L. elongatus* fed constantly throughout the year, while *L. grandis* did not feed in June. Throughout the year, both goby species fed more during the nighttime than the daytime.

**Fig 6.**
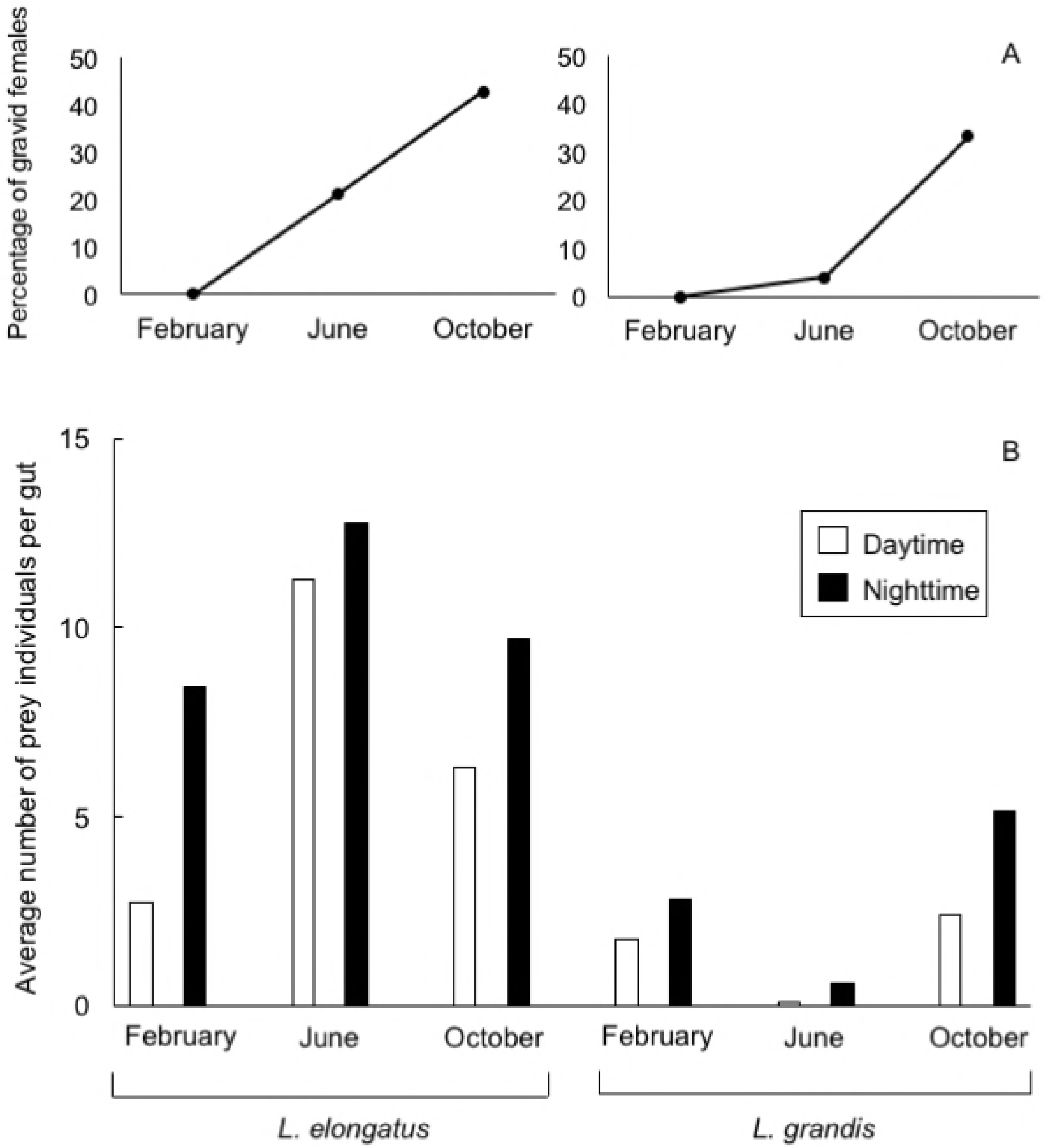
Nonmetric multidimensional scaling ordination of gobies’ prey assemblages. Black and red symbols denote *Luciogobius elongatus* and *L. grandis*, respectively; open and solid symbols denote daytime and nighttime samples, respectively; triangle, circle, and diamond denote February, June, and October, respectively. X marks denote prey categories. Arrows show vectors. The fishes whose guts contained only one individual were excluded.

### Interstitial organism community

Pebbly sediments of *Luciogobius* gobies’ microhabitats harbored diverse and abundant interstitial organisms (Table 2). The great majority of the interstitial organisms were minute meiobenthos under 1 mm in length. The most abundant organisms were harpacticoids, while foraminiferans, nematodes, gastropods, ostracods, amphipods, and isopods were also frequently observed. The community structure was roughly similar between the microhabitats of the two species and between daytime and nighttime, but varied among seasons. Specifically, in October amphipods decreased and foraminiferans increased, and in February flabelliferans increased. Turbellarians, bivalves, mites, and collembolans were observed in sediment but not in gut contents.

**Table 2.** Interstitial organisms found in pebbly sediment inhabited by *Luciogobius elongatus* and *L. grandis*, including the numerical proportions (%) of each organism group obtained during the daytime and nighttime.

| Phylum | *L. elongatus* |  | *L. grandis* |  |
| --- | --- | --- | --- | --- |
| Subphylum |  |  |  |  |
| Class |  |  |  |  |
| Subclass | Daytime | Nighttime | Daytime | Nighttime |
| Order |  |  |  |  |
| Suborder |  |  |  |  |
| Infraorder | n (3918) | n (1187) | n (4500) | n (666) |
| Sarcomastigophora |  |  |  |  |
| Rhizopodia |  |  |  |  |
| Foraminiferida | 10.1 | 9.8 | 1.0 | 1.2 |
| Platyhelminthes | 0.4 | 2.3 | 0.7 | 5.4 |
| Nematoda | 5.4 | 11.4 | 0.3 | 0.3 |
| Mollusca |  |  |  |  |
| Bivalvia | 0.4 | 0.3 | 0.1 | 0.2 |
| Gastropoda | 7.0 | 11.0 | 0.4 | 0.5 |
| Annelida | 1.8 | 1.1 | 3.6 | 5.1 |
| Arthropoda |  |  |  |  |
| Chelicerata |  |  |  |  |
| Arachnida |  |  |  |  |
| Acarina | 0.3 | 0.1 | 0.0 | 0.5 |
| Crustacea |  |  |  |  |
| Ostracoda | 3.6 | 4.5 | 1.4 | 4.4 |
| Maxillopoda |  |  |  |  |
| Copepoda |  |  |  |  |
| Harpacticoida | 63.0 | 51.8 | 87.3 | 67.4 |
| Malacostraca |  |  |  |  |
| Eumalacostraca |  |  |  |  |
| Mysidacea | 0.0 | 0.0 | 0.0 | 0.0 |
| Amphipoda | 6.1 | 5.1 | 2.5 | 8.0 |
| Isopoda |  |  |  |  |
| Asellota | 0.1 | 0.1 | 0.0 | 0.3 |
| Flabellifera | 1.7 | 2.2 | 1.4 | 6.2 |
| Tanaidacea | 0.1 | 0.0 | 0.0 | 0.2 |
| Cumacea | 0.0 | 0.1 | 0.0 | 0.0 |
| Decapoda |  |  |  |  |
| Pleocyemata |  |  |  |  |
| Brachyura | 0.0 | 0.1 | 0.0 | 0.0 |
| Insecta |  |  |  |  |
| Collembola | 0.1 | 0.3 | 1.3 | 0.6 |

### Comparison between prey assemblages and interstitial organism communities

The prey assemblages were roughly similar to the interstitial organism communities, which consisted of harpacticoids, flabelliferan and asellotan isopods, amphipods, brachyurans, ostracods, caecid gastropods, and annelids (Figs 7, 8). The prey assemblages were dominated in number by harpacticoids, which were the most abundant interstitial organisms. While annelids, nematodes, and foraminiferans were common in the sediment, they were rare in gobies’ guts.

**Fig 7.**
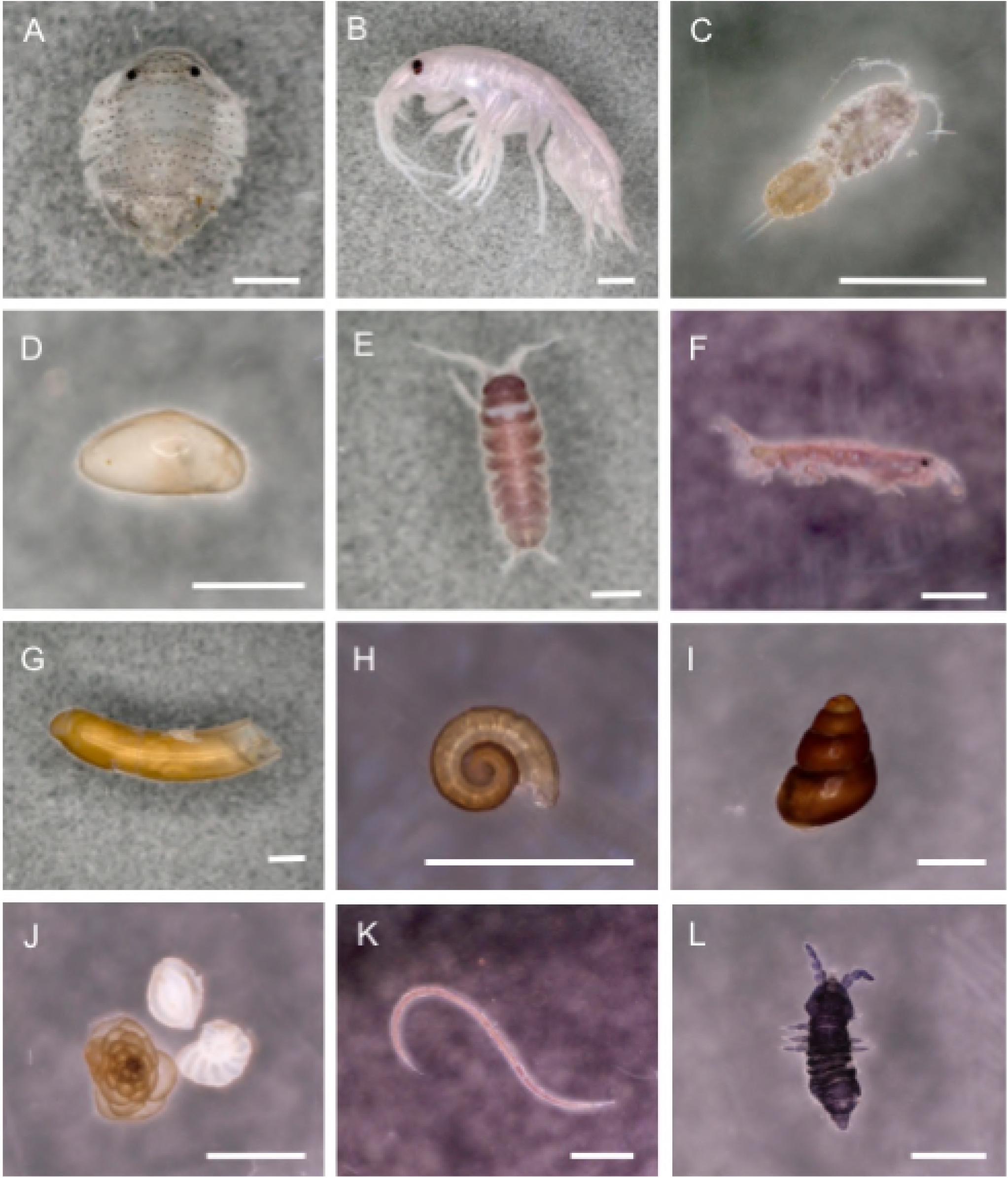
Seasonal and diurnal comparisons of the prey assemblage in the gut of *Luciogobius elongatus* with the meiobenthic community in their microhabitat: (A–C) daytime; (D–F) nighttime; (A, D) February; (B, E) June; (C, F) October.

**Fig 8.**
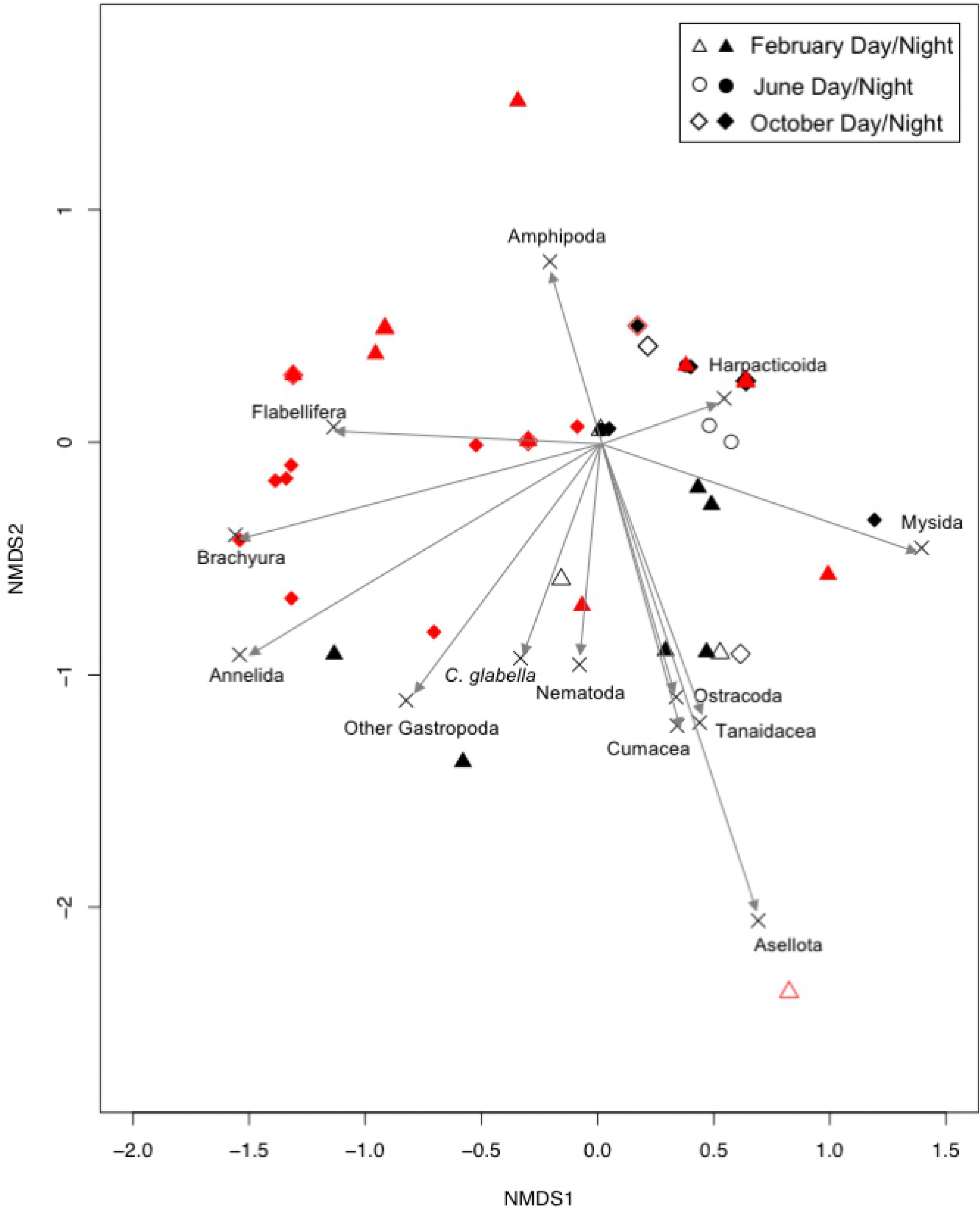
Seasonal and diurnal comparisons of the prey assemblage in the gut of *Luciogobius grandis* with the meiobenthic community in their microhabitat: (A–C) daytime; (D–F) nighttime; (A, D) February; (B, E) June; (C, F) October.

Irrespective of the rough similarity between the prey assemblages and the interstitial organism communities, the proportions of some groups differed. In *L. elongatus*, the proportions of harpactioids in gut were greater than those in sediment, especially in June and October and in both daytime and nighttime (Fig 7), suggesting selective predation of harpacticoids. In *L. grandis*, the proportions of flabelliferan isopods and amphipods in gut were greater than in sediment in both daytime and nighttime (Fig 8), suggesting that they were also selectively predated. Because *L. grandis* gut contents in June were frequently empty, the proportions of amphipods and flabelliferan isopods were 100%, respectively (Fig 8).

The numerical dominance of harpacticoids in diets does not necessary indicate nutritional importance, because harpacticoids are minute. The size distribution of interstitial organisms collected at the gobies’ habitats (Fig 9) showed that flabelliferan isopods were exceptionally large among the interstitial organisms. Thus, the prey assemblages of the two goby species were dominated in biomass by isopods (especially suborder Flabellifera), which constituted less than 50% in number but 60–80% in terms of volume in the diet (Fig 10). Numerical and volumetric comparisons between prey assemblages and interstitial organism communities (Fig 10) showed that the numerical dominance of harpacticoids in the *L. elongatus* gut was reversed in the volumetric comparison, and that the numerical dominance of isopods in the *L. grandis* gut was reinforced in the volumetric comparison.

**Fig 9.**
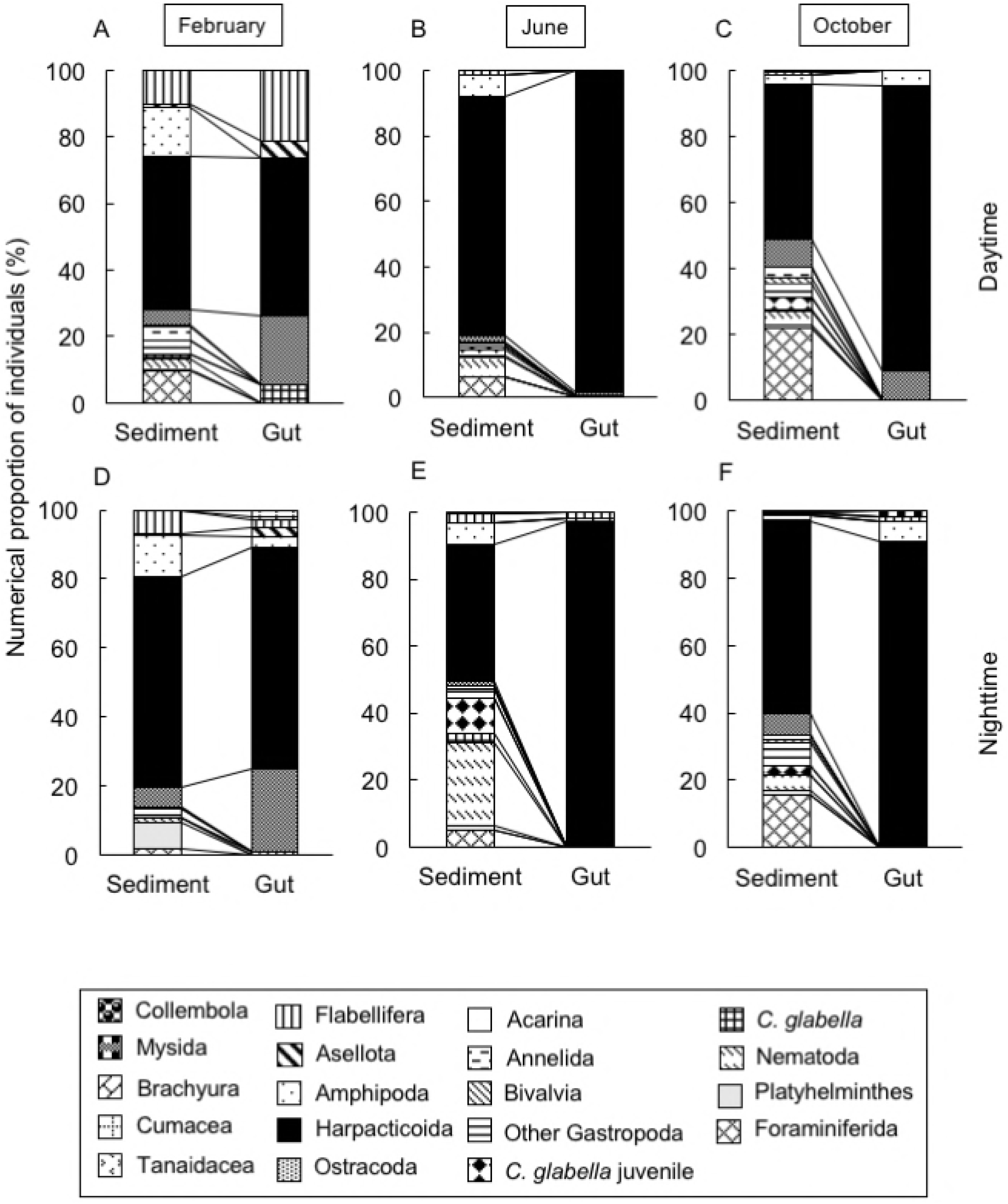
Mean biomass of the main interstitial meiobenthos groups collected from the gobies’ microhabitats. Bar = standard deviation.

**Fig 10.**
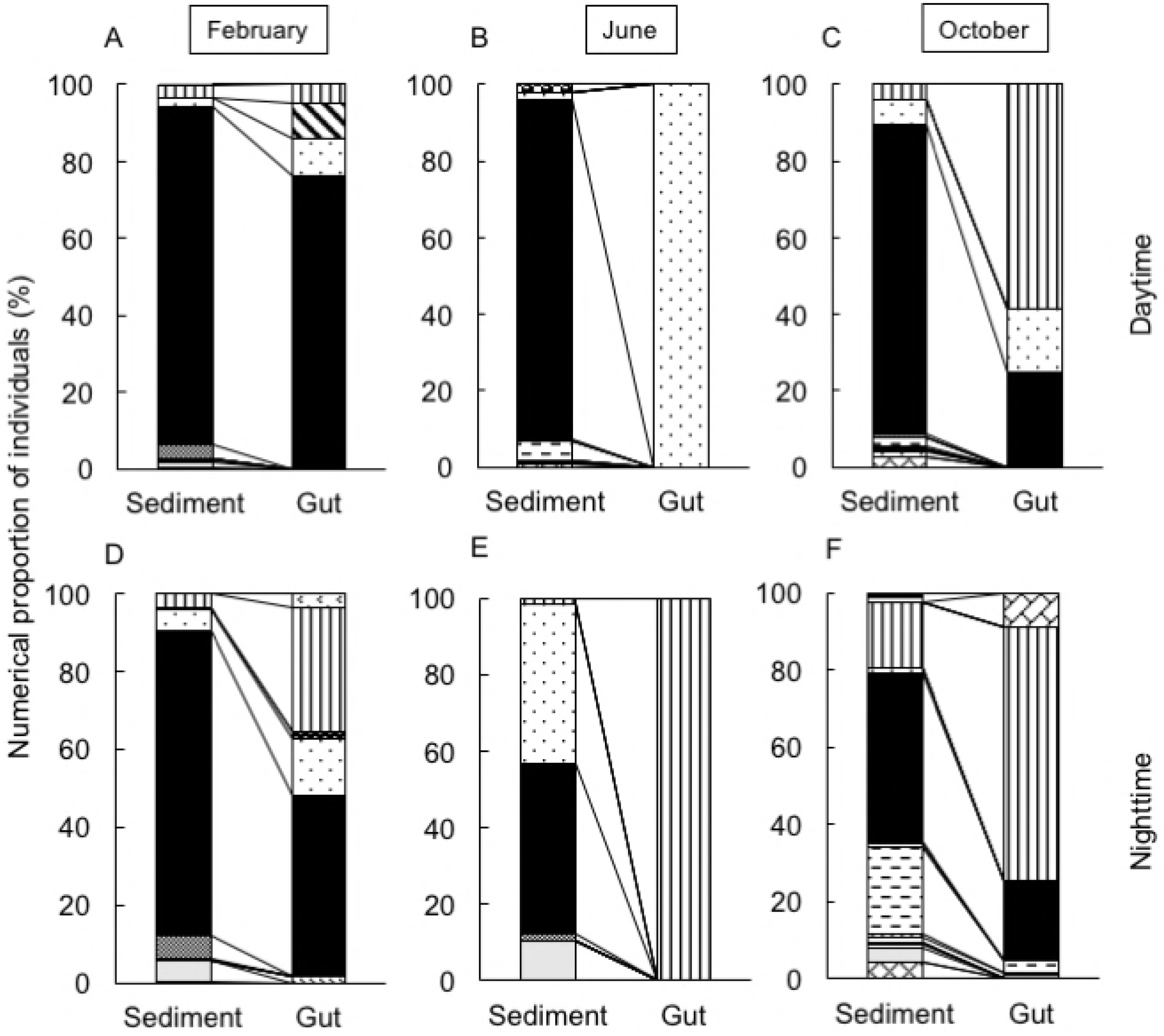
Numerical and volumetric comparisons of the prey assemblage in guts of the two *Luciogobius* species with the meiobenthic communities in their respective microhabitats: (A, B) numerical comparison; (C, D) volumetric comparison; (A, C) *L. elongatus*; (B, D) *L. grandis*.

## Discussion

Pebbly sediments of gravelly coasts in the Japanese Archipelago are unique habitats inhabited by diverse *Luciogobius* gobies and diverse interstitial organisms. This is the first paper reporting the interstitial organism communities in such habitats. The interstitial organism communities were dominated by minute arthropods such as harpacticoids, flabelliferan and asellotan isopods, amphipods, and ostracods (Table 2), presenting a marked contrast with the communities observed in sandy sediments on sandy beaches, where nematodes, turbellarians, annelids, and tardigrades are dominant [10–15] (Fig 11). The interstitial organism community in shelly gravel sediment in Britain was intermediate between our pebbly sediments and the sandy sediments [15]. Interstitial organisms in pebbly sediment are larger and more elastic or more armored than those in sandy sediment, partly because pebbly sediment harbors larger interspaces and partly because pebbly sediment is stirred more violently by strong waves. The most outstanding difference in interstitial faunae between pebbly and sandy sediments is the presence of interstitial vertebrates in the former, e.g., *Luciogobius* gobies with highly extended elastic bodies.

**Fig 11.**
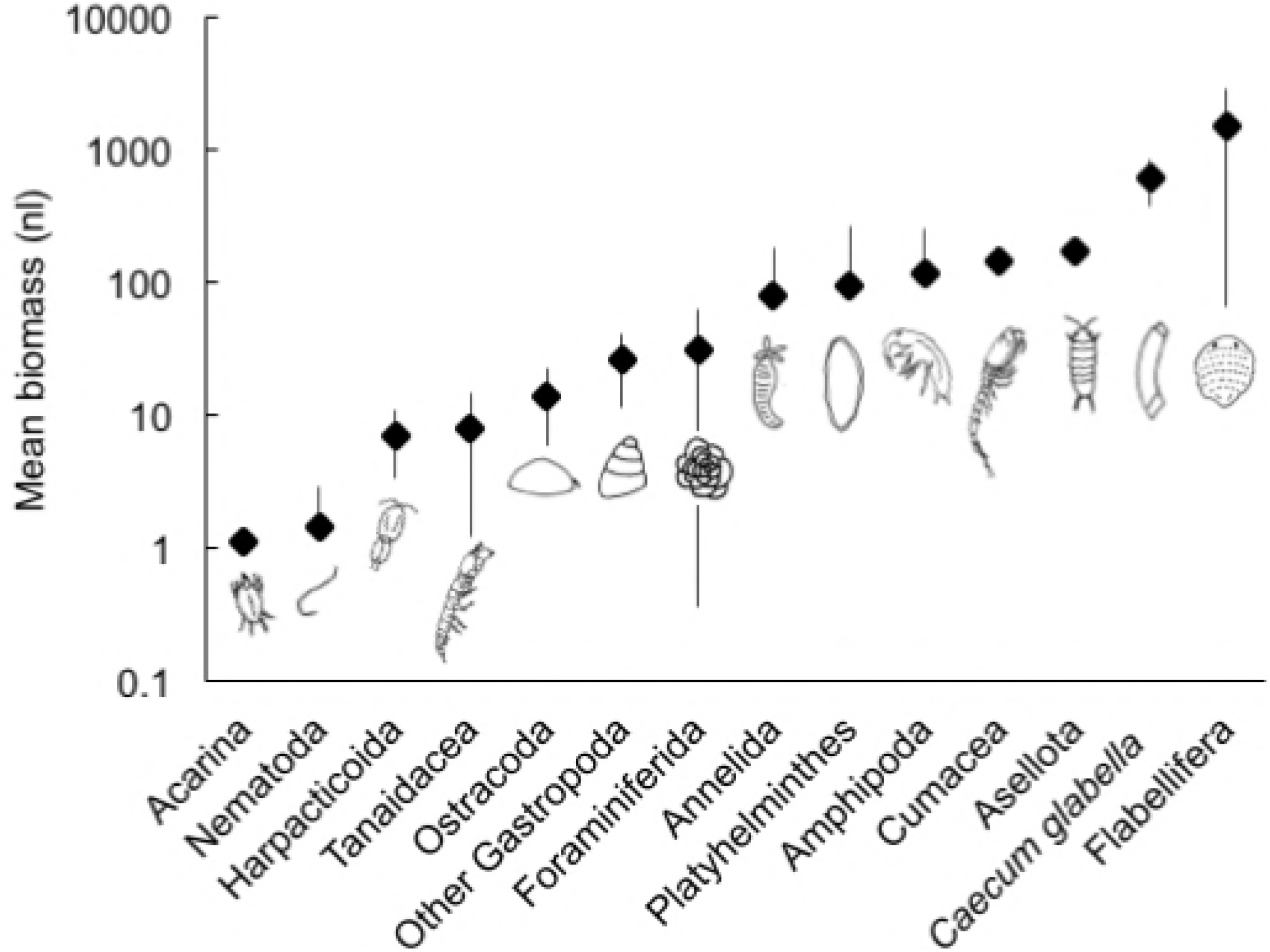
Comparisons of community structure of interstitial organisms among sandy, shelly, and pebbly beaches. The data sources are as follows: a, Kotwicki (2005); b, Ito (1984); c, Williams (1972); d, this study.

Our data on *Luciogobius* goby gut contents and interstitial organism communities within their habitats showed that dietary prey assemblages and interstitial communities were largely similar (Figs 7, 8). The prey assemblages comprised mainly harpacticoids, flabelliferan and asellotan isopods, ostracods, amphipods, and caecid gastropods (Table 1), all of which were small (less than 3 mm in length; the great majority less than 1 mm), armed or unarmed, and swim or crawl in interstitial areas between pebbles. Irrespective of the rough similarity between prey assemblages and interstitial organism communities, *L. elongatus* exhibited a preference for harpacticoids, ostracods, and caecid gastropods, while *L. grandis* exhibited a preference for flabelliferan isopods, amphipods, and brachyurans (Figs 7, 8). The guts contained almost no sand particles, but contained many shelled organisms such as ostracods and caecid gastropods, suggesting that the gobies recognize shelled organisms as available prey.

The average number of gut items differed between goby species (Fig 4B). *L. grandis* fed more in October than February and June. The proportion of flabelliferans in sediment increased in October (Fig 8), which may explain the pattern in *L. grandis* gut contents.

Although harpacticoids were most abundant among prey items, flabelliferan isopods were predominant based on volume (Fig 10), suggesting that the interstitial flabelliferan isopods were staple food resources for the gobies. The scarcity of nematodes and polychaetes among prey items for *Luciogobius* gobies is striking, because sand-burrowing sand darts, *Kraemeria cunicularia*, feed on nematodes and polychaetes in addition to harpacticoids [16]. The scarcity of nematodes and polychaetes in the *Luciogobius* gobies’ gut contents is thought to result from scarcity of these organisms in pebbly sediment compared with sandy sediment. Although sand darts burrow into sand during the day and swim in the water column at night to catch plankton [16], the two *Luciogobius* gobies stayed in the pebbly interstitial environment throughout the day and throughout tidal levels.

The prey assemblages of the two interstitial *Luciogobius* species were roughly similar, suggesting that their feeding behavior is also similar. This result was unexpected, because *L. grandis* is larger and thicker than *L. elongatus*. The dietary similarity of the two goby species suggests that there must be competition for prey between the species, and this may be another reason why the two goby inhabit different microenvironments (i.e., direct behavioral interference causes microhabitat segregation).

Irrespective of the dietary similarity between the two goby species, a preference for harpacticoids was detected in *L. elongatus* (Fig 7) and a preference for flabelliferan isopods was observed in *L. grandis* (Fig 8). Selective feeding on meiobenthos may facilitate the coexistence of several interstitial goby species in pebbly sediment.

The prey assemblages of the gobies were largely similar between night and day (Figs 7, 8). In general, most gobies are visual feeders and feed during the daytime [17–21]. In this survey, the night sampling was conducted at midnight, and fresh prey were found in the guts of *Luciogobius* gobies collected at night. These results suggest that they feed upon interstitial organisms even at night, and that they can find and collect prey in the dark using olfactory and/or tactile senses. Because pebbly interstitial zones are unique habitats where it is dark and visibility is highly restricted, inhabitants have a limited need for sight. Therefore, in interstitial *Luciogobius* gobies, eyes have become vestigial, and various sensory organs around the mouth have developed [5].

Diverse goby species inhabit diverse habitats such as tide pools, tidal flats, estuaries, rocky reefs, coral reefs, and sandy beaches, and their prey are also diverse, e.g., algae, plankton, benthic animals (especially small crustaceans and polychaetes), and even fishes [22–26]. This is the first report of the diet of interstitial gobies, and the *Luciogobius* species studied are the first fishes shown to depend exclusively on interstitial organisms. The prey assemblages of these interstitial *Luciogobius* species contrast with that of lapidicolous *L. guttatus*, which is dominated in number by harpacticoids and in dry weight by juvenile crabs [7]. At our study site in Shirahama, a crab species *Cyclograpsus pumilio* (Varunidae: Decapoda) [27] inhabits pebbly sediment. Although there were no adult *C. pumilio* in the gobies’ guts, very few juvenile crabs were found in the *L. grandis* gut. Thus, the juvenile crabs inhabiting pebbly sediment are also a potential food resource for interstitial *Luciogobius* gobies. The finding that interstitial gobies feed exclusively on interstitial organisms provokes a new question: namely, how do gobies hunt for prey within their interstitial microhabitats, where visibility is restricted and pebbles are continuously stirred by waves?

## Acknowledgements

We are grateful to all the staff of the Seto Marine Biological Laboratory and Shirahama Aquarium for supporting our survey; and Luna Yamamori of Kyoto University for field assistance.

The English in this document has been checked by at least two professional editors, both native speakers of English. For a certificate, please see: http://www.textcheck.com/certificate/vL01DS

